# Methods for Quasispecies Single Observations Comparative Analysis

**DOI:** 10.1101/2024.12.30.630765

**Authors:** Josep Gregori

## Abstract

The study of viral quasispecies structure and diversity poses distinct challenges when comparing samples, particularly in scenarios involving single observations from individual biosamples collected at different time points or under varying conditions. In such cases, each biosample is summarized by a quasispecies haplotype rank-abundance vector, rendering traditional statistical methods inapplicable due to the lack of replicates. Quasispecies structure indicators are functions of multinomial parameters, whose asymptotic normality is guaranteed by the continuous differentiability of these indicator functions and sufficiently large sample sizes. Accurate variance estimation for these indicators is critical for reliable inference. This study contrasts three approaches for variance estimation: analytical first-order approximations via the Delta method, and variance estimates obtained through Bootstrap resampling (with replacement, infinite population), and through resampling without replacement using the random delete-*d* jackknife (finite sample). Exact binomial variances of selected proportions are used as benchmarks to calibrate the resampling methods. We analyze the intrinsic sampling properties of each quasispecies indicator using high-depth next-generation sequencing data from a hepatitis C virus (HCV) cell culture experiment. Our results show that while a subset of indicators yield comparable and close variance estimates across methods, metrics highly sensitive to fluctuations in rare haplotype fractions exhibit substantial divergence. Limitations inherent to the resampling methods are discussed. The analytic approach and tests based on normality are finally recommended.

## 1 Introduction

A quasispecies is the complex and dynamic population of closely related but genetically diverse viral variants that arises during RNA virus infection. This population structure arises due to the high mutation rates of RNA viruses, which can introduce multiple errors during genome replication. The quasispecies consists of a mutant spectrum or cloud of viral genomes that are continuously generated and subject to selection within the host [1–3].

A quasispecies observation consists in sequencing to high depth a quasispecies sample taken from a biosample infected with a RNA virus, and it is represented by a set of genomes (haplotypes) and observed frequencies, in the form of NGS sequencing read counts. We focus on quasispecies structure indicators and diversity indices which may be calculated directly from the rankabundance distribution, that is the ordered vector of haplotype NGS read counts.

In the study of quasispecies structure and diversity, we are often confronted with a situation where we need to compare single quasispecies observations from single biosamples, possibly taken at two different infection or treatment time points. This situation presents statistic challenges, because traditional inferencial tests are not applicable when comparing diversity indices or structure indicators of single quasispecies observations. A useful test must be based on estimates of expected diversity and its variability for each of the quasispecies to be compared.

Analytical approximations derived by the Delta method are available [4–7]. Alternatives are based on resampling to generate empirical distributions from which estimate standard errors. A sound and commonly used resampling method to estimate variances is the bootstrap [8–10], At each bootstrap cycle, a sample is drawn with replacement from the original data, maintaining the same sample size. Diversity indices are computed for each bootstrap sample, and after B iterations, estimates and corresponding variances are derived from the mean and variance of these bootstrap replicates. The standard bootstrap method implicitly assumes an infinite population sampling model. This means that when resampling with replacement from the observed data, the bootstrap treats the data as if drawn from an infinitely large superpopulation. This assumption, often unstated, is inherent in the bootstrap procedure [8,11,12]. Sampling with replacement in this finite population induces a known limitation commonly referred to as the 0.632 problem. In each bootstrap sample, on average, only about 63.2% of the original reads appear as unique values; approximately 36.8% of the reads are duplicates, causing about 36.8% of the original reads to remain unobserved in any given bootstrap replicate [12,13]. This fraction follows from the limit as the sample size *n* tends to infinity of the probability of observing a read in a bootstrap sample, computed as 1 − (1 − 1*/n*)^*n*^, which converges to 1 ^−^ 1*/e* = 0.6321. This limitation can possibly bias estimates of diversity indices sensitive to the presence of very rare haplotypes. Indices such as the total number of haplotypes, Shannon entropy, and the Hill number at *q* = 1 can be particularly affected [9,13–16].

Note that any approach based on parametric multinomial resampling is subject to the same limitations: resampling from a multinomial distribution given its parameters and a sample size is equivalent to resampling from an infinite population. In other words, it corresponds to sampling with replacement from a sample where the probabilities are fixed and the draws are mutually independent.

In this context, an alternative to the bootstrap is the jackknife [17]. While similar to the bootstrap in that it involves resampling, the key difference is that the jackknife samples without replacement, which aligns with the method used in rarefaction [13–15] to normalize sample sizes. However, it is well known that the classical delete-1 jackknife estimator performs poorly with non-smooth estimators—those that are sensitive to one or a few values in the dataset, such as the median or other empirical quantiles. This makes it particularly unsuitable for computing confidence intervals. It has been shown that the delete-d Jackknife [8,18], where 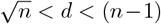, is a consistent estimator [8,19,20] and overcomes most of the limitations in the delete-1 jackknife. This method has a similar form to the delete-1 estimator but incorporates a different normalizing constant, accounting for the number of distinct sets with *d* items deleted from the full set of *n*,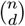. However, as the complexity of the computations increases rapidly with larger *d* -especially for large *n*-the estimator may be evaluated for a subset of all possible 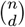 sets [8,20–22].

Shao’s [20–22] formulation of the delete-d jackknife variance estimator corrects the bias arising from two sources when estimating the variance of an estimator based on a full sample of size *n*: 1-the reduced size *n* − *d* of jackknife subsamples, and 2-the positive covariances between jackknife replicates caused by overlapping subsets. The variance computed from jackknife replicates, each based on a subsample of size *n* − *d*, naturally overestimates the estimator’s true variance due to smaller sample size. Furthermore, because deleted subsets overlap, the jackknife replicates are correlated, which reduces their effective variability. Shao showed, using combinatorial arguments on inclusion patterns and covariance structures of observations across subsamples, that these two effects combine to produce a multiplicative correction factor of (*n* −*d*)*/d*. Multiplying the empirical variance of the delete-d jackknife replicates by this factor yields a consistent and unbiased estimator for the variance of the estimator computed from the full sample. In this work, we employ a random delete-d jackknife, in which a fixed fraction of reads is randomly removed across a large number of resampling cycles. This scheme is closely related to repeated rarefaction to a lower sample size, subsampling without replacement. Note that in our context *n* is in the order of 10^5^, *d* is chosen to be a 10%, 10^4^, and the number of jackknife resamplings 10^3^.

We present in the Results section a comparison of exact variances for binomial proportions, analytic variance approximations for functions of the rank-abundance vector obtained using expressions derived via the delta method, variance estimates obtained from 5000 bootstrap iterations, and estimates derived from 5000 resampling cycles of the random delete-d jackknife. The effect of the deleted fraction *d* on the resulting variance estimates is also analyzed. Selected quasispecies fractions serve as benchmarks to calibrate the estimated variances by resampling methods against Bernouilli exact values.

From the literature, a general summary can be extracted [19,23–33]:

- Bootstrap: Superior for small samples, nonlinear estimators, and non-normal errors due to its ability to approximate the entire sampling distribution. However, it can be less efficient computationally than other methods when parametric assumptions hold and sample sizes are large.
- Jackknife: Highly efficient for linear and large-sample problems, often producing unbiased variance estimates but with higher variability than the bootstrap in certain cases.
- Delta method: Analytical and computationally efficient; reliable when the estimator is differentiable and asymptotically normal. Best suited for smooth functions where local linearity holds near the true parameter value.

In quasispecies populations with complex structure, where many haplotypes are represented by few sequencing reads—often occurring at very low frequencies—the typical assumptions underlying resampling-based variance estimation may be violated. In such scenarios, the bootstrap tends to underestimate the true number of haplotypes, failing to capture highly rare variants effectively [13–15]. The jackknife estimator also generally underestimates haplotype richness, though its bias is typically smaller and depends on the proportion of data removed in each resample. Biases in both methods directly impact the *RLE*_*q*_ indicators, and further distortions arise due to inaccurate frequency estimates for the rarest haplotypes present in resampled datasets. Nevertheless, delta method variance formulas for *RLE*_*q*_ treat *H*, the number of haplotypes in the entire sample, as a known non-random parameter. This same assumption, for comparative reasons, is adopted in the bootstrap and jackknife calculations: *H* is taken as the cardinality of the haplotypes observed in the complete sample, even though some haplotypes may be absent from specific resampled datasets.

The data used in this analysis belongs to a study of in-vitro HCV quasispecies evolution, with samples at baseline and at different passages up to 200, representing the quasispecies evolution in a controlled non-coevolving cellular environment free of external pressures, for 700 days [34,35].

This study aims to systematically compare exact variances for proportions, and variance estimates obtained from analytical expressions derived via the delta method with those estimated by the two mentioned resampling approaches. By evaluating the agreement and discrepancies between these methods, we seek to identify potential sources of deviation and inform the choice of variance estimation techniques for complex estimators in finite samples of complex populations. Here, variances are treated as independent of sample size, based on the relationship 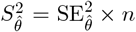, where 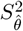 is the sample variance, 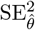 is the squared standard error of the estimator 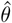, and *n* is the sample size. This distinction is particularly important when calculating effect sizes, which provide metrics that are independent of sample size, enabling meaningful comparisons among quasispecies populations. Such effect size measures offer insights beyond mere statistical significance by quantifying the magnitude of observed differences.

## 2 Methods

### 2.1 Quasispecies maturity indicators

Table 1 shows the selected set of quasispecies structure indicators designed to characterize the maturation and evolutionary state of a quasispecies [4,36,37]. This progression traces the path from a founding quasispecies, A, dominated by a highly abundant master genome with a peaked fitness landscape, toward a hypothetical endpoint, Z, characterized by a flat fitness landscape. At this final stage, a large number of genomes exist at low and approximately equal abundances, in dynamic equilibrium, with no dominant variants [36,37].

**Table 1:** Selected maturity indicators. QFF: Quasispecies Fitness Fractions [38]. QsE: Quasis-pecies evenness indicators [36]. TopE: Evenness among top haplotypes [4,36].

| Indicator $f$ | Type | Description | $f(A)$ | $f(Z)$ |
| --- | --- | --- | --- | --- |
| TopN | QFF | Fraction of reads for top N hpl. | 1 | 0 |
| Master | QFF | Dominant haplotype frequency | 1 | 0 |
| Rare1 | QFF | Fraction of reads for hpl $\leq 1\%$ | 0 | 1 |
| Rare2 | QFF | Fraction of reads for hpl $\leq 0.1\%$ | 0 | 1 |
| $RLE_1$ | QsE | Relative logarithmic evenness at $q = 1$ | 0 | 1 |
| $RLE_2$ | QsE | Relative logarithmic evenness at $q = 2$ | 0 | 1 |
| $RLE_{\infty}$ | QsE | Relative logarithmic evenness at $q = \infty$ | 0 | 1 |
| $R_k$ | TopE | Evenness among top k haplotypes | 0 | 1 |

These indicators are functions of the haplotype rank-abundance vector:

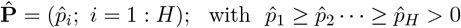

where *H* is the cardinality of the observed haplotypes, the dimension of the vector 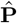.

The zeroes for *TopN* and *Master*, in state Z, should be interpreted as an infinitesimal frequency *ϵ*, that is *ϵ >* 0 and *ϵ <* 1*/n* for a big positive integer *n*, rather than strictly zero. In biochemical terms this *ϵ* may be interpreted as one to a few or several molecules, in the context of a high viral titer. Similarly, the zeroes for Rare1 and Rare2, in state A, should be interpreted as a result of several variants other than the master haplotype, at infinitesimal frequencies, *ϵ*. However, we represent *ϵ* formally as 0.

Quasispecies fitness fractions (QFF) are binomial proportions, or aggregates of multinomial proportions that can be treated as binomial proportions, computed from the estimated haplotype rank-abundance vector 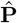. *QFF* indicators summarize key fractions in the composition of the quasispecies, such as the proportions of master, emerging, low fitness, and very low fitness haplotypes, and provide insight into the population’s structure.

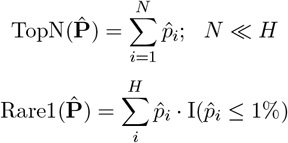

where I(·) is the indicator function, which returns 1 if the condition is fulfilled and 0 otherwise.

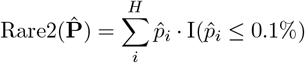

Relative logarithmic evenness (RLE) indicators of order *q, RLE*_*q*_, provide measures of the evenness in the distribution of haplotypes.

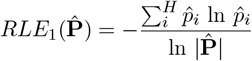

with 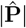 the dimension of the rank-abundance vector, the number of observed haplotypes.

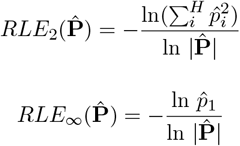

Top haplotypes evenness quantifies the degree of evenness among the most abundant haplotypes, providing insight into how similarly represented they are within the quasispecies population.

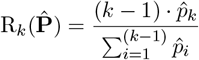

Finally, a global quasispecies maturity score for a given quasispecies with estimated rank-abundance vector, 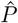, may be computed as the distance from this quasispecies to state A, *d*_*A*_, in the multidimensional space represented by this set of indicators [4].

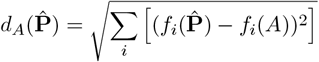

where *f*_*i*_ is the set of maturity indicators. This score may be normalized to the range 0-1 as 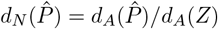. Whereas each indicator refers to a given aspect of quasispecies structure, *d*_*N*_ represents the combined effect of all changes in pushing the quasispecies towards its origin, A, or its fate, Z [4].

### 2.2 Quasispecies fitness fractions, exact variances

Quasispecies fitness fractions -*Master, Top25, Rare1*, and *Rare2* - are sums of haplotype frequencies of designed types, and can be modeled as either aggregates of multinomial parameters or as binomial proportions, allowing exact calculation of their variance.

Let 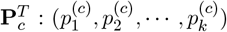, with *k < H*, denote the frequencies of a given set of haplotypes within class *c*. Define their sum as:

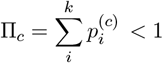

If Π_*c*_ is treated as a binomial parameter (interpreted as the probability of success in a Bernoulli trial), its variance is given by the Bernouilli formula:

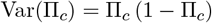

Alternatively, if the components 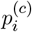 are considered as parameters of a multinomial distribution, they are not independent but negative correlated due to the multinomial constraint. The covariance matrix for these multinomial components, **Σ**_*c*_, is expressed in matrix algebra as:

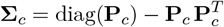

where 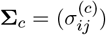 has elements:

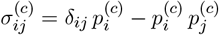

with *δ*_*ij*_ the Kronecker delta function, which equals 1 if *i* = *j* and 0 otherwise.

The total variance of the sum Π_*c*_ can be computed as the sum over all covariance terms:

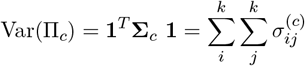

where **1** is the all-ones vector with dimension *k*.

It can be shown that the binomial variance equals the variance of the sum derived from the multinomial covariance matrix, both approaches are mathematically equivalent.

Binomial and multinomial parameters are asymptotically normal due to the Central Limit Theorem (CLT), which states that as the sample size *n* increases, the sampling distributions of their estimators approximate normal distributions regardless of the underlying discrete nature. For the binomial parameter *p*, the maximum likelihood estimator 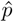 converges in distribution to a normal distribution with mean *p* and variance *p*(1− *p*)*/n*. The multinomial distribution generalizes this to a vector parameter **P** = (*p*_1_, …, *p*_*k*_) where the vector of sample proportions 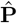 approaches a multivariate normal distribution, centered at **P** with covariance matrix reflecting the multinomial constraints. This asymptotic normality facilitates statistical inference by enabling the use of normal-based confidence intervals and hypothesis tests for proportions and categorical data even when the underlying distributions are discrete.

### 2.3 Variances approximated using the delta method

Quasispecies evenness, *RLE*_*k*_, and top haplotypes evenness indicators, *R*_*k*_ are functions of the estimated haplotype rank abundance vector, 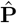, and its variance may be approximated by expressions derived using the delta method. Refer to [4]-and references therein-where the equations for these indicators have been derived. This section provides the elements of the delta method and its use in inference.

The delta method [39–42] provides a statistical foundation for justifying normality in tests involving functions of random variables by leveraging the asymptotic normality of the parameters estimators. When an estimator is asymptotically normal, the delta method allows us to approximate the distribution of smooth (i.e. differentiable) functions of that estimator as also normal. The proposed quasispecies structure indicators are based on the distribution of haplotypes, which follows a multinomial distribution with as many categories and parameters as there are haplotypes. Multinomial proportions estimated from a sample exhibit asymptotically normal behavior,

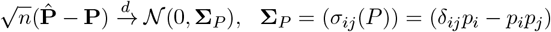

providing the basis for applying the delta method to functions of estimated proportions 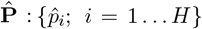, and proportion aggregates, and for conducting tests based on the normal distribution using these indicators. In what follows, let 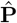 denote the vector of observed haplotype frequencies, sorted in decreasing order of frequency without loss of generality, so that 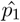 represents the estimated frequency of the master haplotype.

Briefly, if an estimator 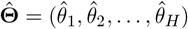 satisfies

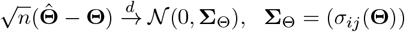

then for a differentiable function *g*, the delta method ensures:

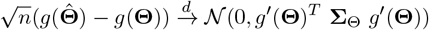

provided that *g*^*′*^(**Θ**)^*T*^ **Σ**_**Θ**_ *g*^*′*^(Θ) *>* 0, being

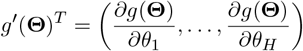

This allows normality tests to be applied to 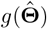, even if *g* is non linear.

The method approximates the standard error of 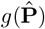 of first order Taylor expansion:

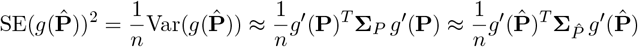

where 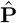 is a vector of multinomial proportions, or an aggregate of them.

The delta approximation is most reliable when the underlying estimator has small variance or large sample size, *g*(**Θ**) is continuously differentiable in a neighborhood of the true value, and the region of interest exhibits local linearity, i.e., higher-order derivatives are small relative to the first derivative in magnitude.

Given the estimates of a quasispecies structure indicator, *g*(·), for two quasispecies being compared, 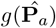 and 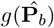, and their standard errors obtained from the delta method, 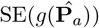 and 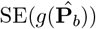, we may compute the z statistic as:

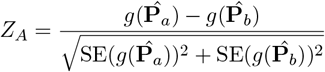

This statistic assesses the significance of the difference between the two quasispecies indicators under the assumption of independent normal errors.

Using the approximate normality, a two-sided p-value may be calculated as

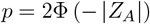

Similarly, a Cohen’s *d*_*A*_ effect size between the two compared conditions may be approximated as:

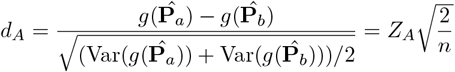

with *n* the normalized sample size of the two quasispecies.

The variance expressions for all proposed quasispecies structure indicators derived using the delta method are provided in [4].

Note that, for consistency across the studied methods, the haplotype cardinality is defined as the number of haplotypes observed on the full sample, as if it was known constant. This is the method used in the variance equations derived using the delta method for *RLE*_*q*_ indicators.

### 2.4 Variances estimated by bootstrap

The standard error of a function of a random variable estimated by the bootstrap method [8–11,24,29] is obtained through a resampling approach. The fundamental idea is to:

1-Draw a large number, *B*, of bootstrap samples, with rank-abundance vector 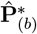, from the original vector of haplotype read counts, 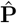, by **sampling with replacement** to the full sample size.

2-Compute the statistic (function) of interest,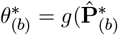, at each iteration *b*, from bootstrap sample, 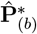.

3-Estimate the standard error of the statistic as the standard deviation of these bootstrap values:

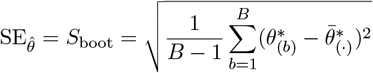

where

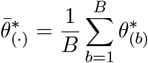

is the mean of the bootstrap values.

This variance estimate approximates the variance of the statistic as if it were computed from the true sampling distribution. The bootstrap variance is conditional on the empirical distribution function of the sample and converges to the true variance as the number of bootstrap samples *B* and the original sample size *n* grow large. It works for any smooth or nonsmooth function of the sample, making the method very versatile in practice.

### 2.5 Variances estimated by random delete-d jackknife

For a quasispecies represented by haplotype read counts summing to *n*, we remove a random fraction *f* of reads (*d* = ⌈ *n* × *f* ⌉) at each iteration [8,20–22]. To minimize correlations across replicates, the reads are first randomly shuffled. A fraction *f* = 0.10 (10%) was chosen as a compromise between perturbation and stability, providing consistent variance estimation across resampling cycles.

At each iteration, *j*, after randomly removing *d* reads from the original sample, we obtain a rank-abundance vector 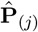, from which we calculate the statistic of interest, 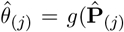. After *R* delete-d jackknife iterations the standard error of the indicator estimate,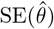, is calculated following Shao’s formulation [19–22] as:

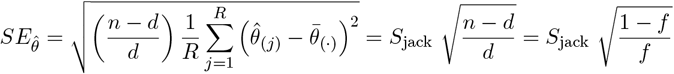

where

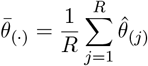

and *d* = ⌈*n* × *f* ⌉, with *f* the fraction of reads to remove in each jackknife resampling iteration.

The factor (*n* − *d*)*/d* corrects for the smaller sample size *d* − *n*, and the covariances between replicates caused by overlapping subsets. Multiplying the empirical variance of the delete-d jackknife replicates by this factor yields a consistent and unbiased estimator for the variance of the estimator computed from the full sample [19–22].

The original standard deviation of the indicator,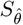, which may be used in the computation of effect sizes, is obtained from the standard error in the usual way:

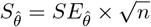

where *n* is the sample size in number of reads.

### 2.6 Overdispersion in resampling

Quasispecies fitness fractions indicators (QFF) represent proportions that can be modeled as binomial parameters, since each sampled read either belongs to a specific *QFF* category or it does not. Overdispersion in sampling from a binomial distribution (with replacement, infinite population) or a hypergeometric distribution (without replacement, finite population) occurs when the observed variability in the data exceeds what the sampling model predicts. Both distributions assume that each trial has an underlying constant probability of success. In our context *success* means that a sampled read belongs to a given quasispecies fitness fraction (i.e. a read of a haplotype with abundance < 0.01%, for fraction *Rare2*).

Overdispersion arises if these assumptions are violated. For example, the underlying probability of success may vary across trials (heterogeneity). This leads to greater variability in the number of successes than the sampling model anticipates.

When the success probability is not constant, the beta-binomial or the beta-hypergeometric distribution provide a natural extension of the basic model. In this compound model, the probability of success is itself a random variable drawn from a beta distribution, representing heterogeneity among samplings.

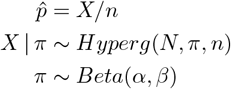

with *X* the number of sampled reads of a given type, *N* the full sample size (population), *n* the resample size, and *α* and *β* the parameters of the beta distribution, and *X/n* provides the estimated proportion. This additional layer introduces an extra dispersion parameter, allowing the variance to be inflated beyond what the basic predicts, thereby capturing overdispersion in the data.

To assess the presence of overdispersion in resampled values a beta distribution is fitted. The parameters of the beta distribution are estimated via maximum likelihood, initializing the optimization with *α* = 1 and 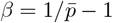, where 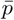 is the mean of the resampled values. These starting values correspond to the simplest beta distribution whose mean matches 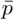.

This beta distribution summarizes the variability in the observed 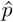 resampled proportions, with a mean value 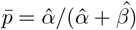 and variance:

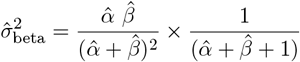

Using these estimates, 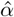 and 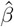, the variance of the beta-binomial distribution with sample size *n* is given by:

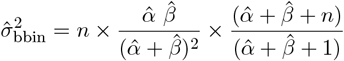

where the last part is the overdispersion factor. Similarly for the beta-hypergeometric model.

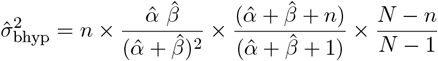

where the last term is the finite population correction factor (FPC).

Hence, the overdispersion relative to the basic model is quantified, using the fitted beta distribution parameters, as:

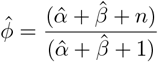

### 2.7 Samples, cells and viruses

The data used in this study correspond to five time points during the evolution of a cell-cultured HCV quasispecies, starting from low diversity and progressively increasing genetic complexity. These five time points represent scenarios of varying complexity, capturing a wide range of values in the quasispecies structure indicators.

HCV RNA expressed from plasmid Jc1FLAG2(p7-nsGluc2A) (genotype 2a) [43] was transfected into Huh-7 Lunet cells and amplified in Huh-7.5 cells to produce the initial virus population HCV p0 [44]. HCV p0 was subjected to 200 serial passages in Huh-7.5 reporter cells in the absence of any drug. Biosamples were taken at different passages: p0, p3, p100, p110 and p200. The populations at passage 100 (HCV p100) and at passage 200 (HCV p200) displayed increased replication in Huh-7.5 reported cells [34,45]. HCV p100 and p200 increased their fitness 2.2-fold relative to HCV p0 [45,46]. The 200 passages are equivalent to about 700 days of continuous evolution in a non-coevolving environment [34,35] free of external pressures. A single NS5A amplicon spanning positions 7523:7864 is deeply sequenced with an Illumina^®^ MySeq instrument for each biosample.

## 3 Results

### 3.1 Samples and NGS coverage

Table 2 lists the quasispecies considered in the tests, and the coverage in number of reads for each strand of the sequenced amplicon.

**Table 2:** Reads per amplicon sample and strand.

| ID | FW | RV |
| --- | --- | --- |
| p0 | 124253 | 120939 |
| p3 | 186398 | 182387 |
| p100 | 179401 | 176168 |
| p110 | 136432 | 133209 |
| p200 | 123807 | 123946 |

In what follows only the fasta files for the forward strands of the amplicon are considered.

### 3.2 Quasispecies maturity indicators

Values for these indicators are calculated directly from the quasispecies haplotype rank-abundance data as observed in the NGS sample, without any normalization for sequencing depth (see Table 3). Our goal is to compare the variances estimated for each quasispecies using the three methods under study; thus, the effective sample sizes are implicitly equivalent, and no normalization is required.

**Table 3:** (A) - Quasispecies maturity indicators.

| ID | nHpl | Master | Top25 | Rare1 | Rare2 | dN |
| --- | --- | --- | --- | --- | --- | --- |
| p0 | 5858 | 0.72884 | 0.80131 | 0.24107 | 0.19969 | 0.18107 |
| p3 | 8325 | 0.73095 | 0.78525 | 0.26905 | 0.18898 | 0.18498 |
| p100 | 11948 | 0.49214 | 0.74294 | 0.33202 | 0.23125 | 0.27198 |
| p110 | 9744 | 0.39303 | 0.75431 | 0.28186 | 0.24154 | 0.29405 |
| p200 | 11490 | 0.43506 | 0.70577 | 0.37739 | 0.25735 | 0.31031 |

**Table 3: (B).**
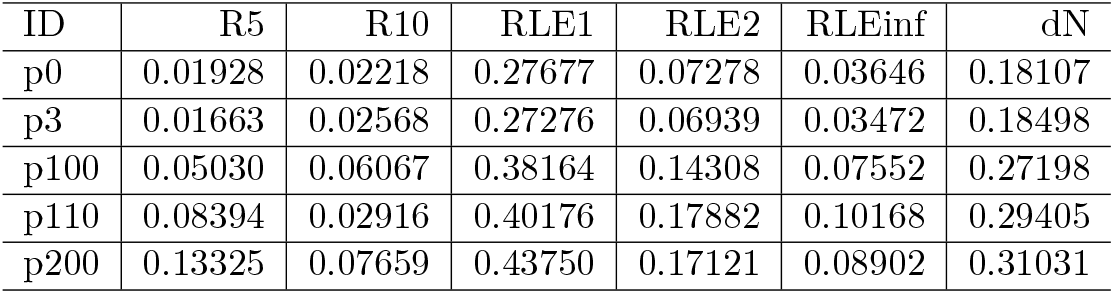
Quasispecies maturity indicators.

| ID | R5 | R10 | RLE1 | RLE2 | RLEinf | dN |
| --- | --- | --- | --- | --- | --- | --- |
| p0 | 0.01928 | 0.02218 | 0.27677 | 0.07278 | 0.03646 | 0.18107 |
| p3 | 0.01663 | 0.02568 | 0.27276 | 0.06939 | 0.03472 | 0.18498 |
| p100 | 0.05030 | 0.06067 | 0.38164 | 0.14308 | 0.07552 | 0.27198 |
| p110 | 0.08394 | 0.02916 | 0.40176 | 0.17882 | 0.10168 | 0.29405 |
| p200 | 0.13325 | 0.07659 | 0.43750 | 0.17121 | 0.08902 | 0.31031 |

Table 3 lists the values of the indicators as computed from the observed read counts.

### 3.3 Exact variances and approximations by the delta method

Table 4 lists the analytic variances -exact for *QFF* indicators, and approximated with delta method for *RLE*_*q*_ and *R*_*k*_ indicators- and computed from the observed vector of haplotype read counts, for each quasispecies and indicator.

**Table 4:** (A) - Quasispecies maturity indicators. Analytic standard deviations.

| ID | Master | Top25 | Rare1 | Rare2 | dN |
| --- | --- | --- | --- | --- | --- |
| p0 | 0.44456 | 0.39902 | 0.42773 | 0.39977 | 0.31634 |
| p3 | 0.44347 | 0.41065 | 0.44347 | 0.39149 | 0.31856 |
| p100 | 0.49994 | 0.43701 | 0.47094 | 0.42163 | 0.34966 |
| p110 | 0.48842 | 0.43050 | 0.44991 | 0.42802 | 0.33234 |
| p200 | 0.49576 | 0.45570 | 0.48473 | 0.43718 | 0.38633 |

**Table 4: (B).** Quasispecies maturity indicators. Analytic standard deviations.

| ID | R5 | R10 | RLE1 | RLE2 | RLEinf | dN |
| --- | --- | --- | --- | --- | --- | --- |
| p0 | 0.31799 | 0.50631 | 0.41546 | 0.14000 | 0.07031 | 0.31634 |
| p3 | 0.29945 | 0.55297 | 0.40870 | 0.13421 | 0.06721 | 0.31856 |
| p100 | 0.55227 | 0.88699 | 0.39744 | 0.18859 | 0.10820 | 0.34966 |
| p110 | 0.70350 | 0.60035 | 0.39217 | 0.19219 | 0.13531 | 0.33234 |
| p200 | 0.95605 | 1.03329 | 0.40151 | 0.21871 | 0.12189 | 0.38633 |

### 3.4 Variances estimated by resampling methods

Table 5 lists the standard deviations estimated by bootstrap, for each quasispecies and indicator.

**Table 5:** (A) - Quasispecies maturity indicators. Boostrap standard deviations.

| ID | Master | Top25 | Rare1 | Rare2 | dN |
| --- | --- | --- | --- | --- | --- |
| p0 | 0.43842 | 0.39416 | 0.42157 | 0.70372 | 0.29898 |
| p3 | 0.44036 | 0.40641 | 0.44036 | 0.71379 | 0.30199 |
| p100 | 0.50099 | 0.43455 | 0.47486 | 0.73002 | 0.30113 |
| p110 | 0.48893 | 0.42810 | 0.46179 | 0.68889 | 0.27862 |
| p200 | 0.50193 | 0.45641 | 0.48501 | 0.75169 | 0.31244 |

**Table 5: (B).** Quasispecies maturity indicators. Bootstrap standard deviations.

| ID | R5 | R10 | RLE1 | RLE2 | RLEinf | dN |
| --- | --- | --- | --- | --- | --- | --- |
| p0 | 0.30750 | 0.29967 | 0.41155 | 0.13806 | 0.06934 | 0.29898 |
| p3 | 0.21977 | 0.31260 | 0.40477 | 0.13326 | 0.06674 | 0.30199 |
| p100 | 0.55900 | 0.72416 | 0.39994 | 0.18933 | 0.10843 | 0.30113 |
| p110 | 0.70695 | 0.53380 | 0.39108 | 0.19182 | 0.13544 | 0.27862 |
| p200 | 0.95270 | 0.81425 | 0.40443 | 0.22112 | 0.12340 | 0.31244 |

Table 6 lists the standard deviations estimated by random delete-d jackknife, for each quasispecies and indicator.

**Table 6:** (A) - Quasispecies maturity indicators. Jackknife standard deviations.

| ID | Master | Top25 | Rare1 | Rare2 | dN |
| --- | --- | --- | --- | --- | --- |
| p0 | 0.44029 | 0.39831 | 0.42621 | 1.17863 | 0.32141 |
| p3 | 0.44830 | 0.40807 | 0.44830 | 1.22467 | 0.32342 |
| p100 | 0.48986 | 0.44651 | 0.47355 | 1.09710 | 0.31235 |
| p110 | 0.48615 | 0.43226 | 0.45137 | 1.11189 | 0.29125 |
| p200 | 0.49892 | 0.45278 | 0.48786 | 1.21479 | 0.32623 |

**Table 6: (B).** Quasispecies maturity indicators. Jackknife standard deviations.

| ID | R5 | R10 | RLE1 | RLE2 | RLEinf | dN |
| --- | --- | --- | --- | --- | --- | --- |
| p0 | 0.31847 | 0.35048 | 0.42679 | 0.13866 | 0.06963 | 0.32141 |
| p3 | 0.24397 | 0.46903 | 0.42192 | 0.13566 | 0.06794 | 0.32342 |
| p100 | 0.55719 | 0.74951 | 0.41362 | 0.18624 | 0.10602 | 0.31235 |
| p110 | 0.71183 | 0.59856 | 0.40984 | 0.19185 | 0.13468 | 0.29125 |
| p200 | 0.94940 | 1.03246 | 0.41887 | 0.22043 | 0.12266 | 0.32623 |

### 3.5 Comparative between methods

#### 3.5.1 Resampled mean values vs observed

A comparative visualization of the quasispecies indicator values calculated directly from the observed haplotype frequencies on the full sample (Observed) and from the mean of resampled values obtained by bootstrap (Boot) and jackknife (Jack) is shown in Figure 1. The corresponding relative differences between the observed and resampled estimates are listed in Tables 7 and 8, and plotted in Figure 2.

**Table 7:** Relative differences in estimated indicator values. (bootstrap-observed)/observed x 100.

| ID | Top25 | Master | Rare1 | Rare2 | R5 | R10 | RLE1 | RLE2 | RLEinf | dN |
| --- | --- | --- | --- | --- | --- | --- | --- | --- | --- | --- |
| p0 | 0.027 | -0.003 | 0.008 | -0.687 | -0.047 | -0.172 | -1.098 | 0.010 | 0.010 | -0.387 |
| p3 | 0.035 | 0.000 | -0.001 | 0.652 | 1.488 | 0.126 | -1.025 | -0.001 | -0.001 | -0.188 |
| p100 | 0.009 | 0.002 | -0.006 | -0.134 | -0.035 | -2.013 | -1.052 | -0.005 | -0.003 | -0.255 |
| p110 | 0.022 | -0.001 | -0.004 | 0.052 | 0.011 | -0.344 | -1.090 | -0.001 | 0.002 | -0.226 |
| p200 | 0.022 | 0.004 | 0.000 | 0.012 | -0.020 | 0.575 | -1.282 | -0.004 | -0.004 | -0.283 |

**Table 8:**
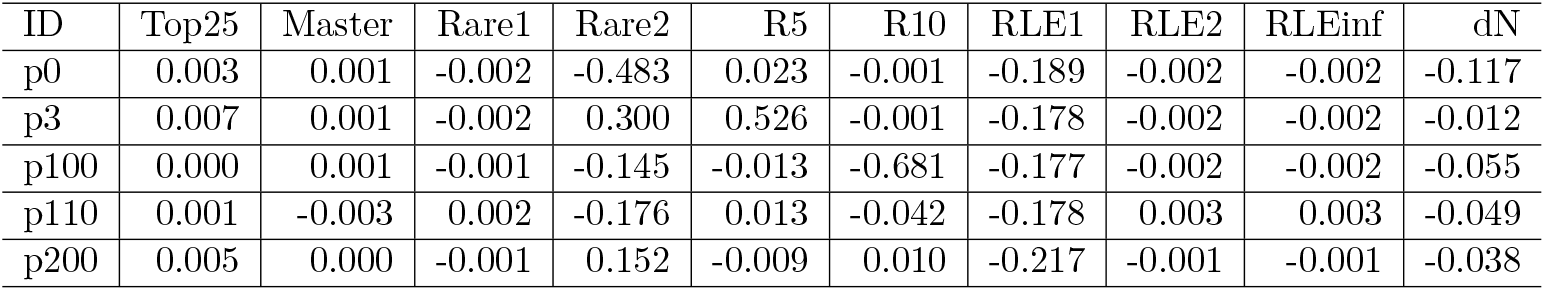
Relative differences in estimated indicator values. (jackknife-observed)/observed x 100.

**Table 9:**
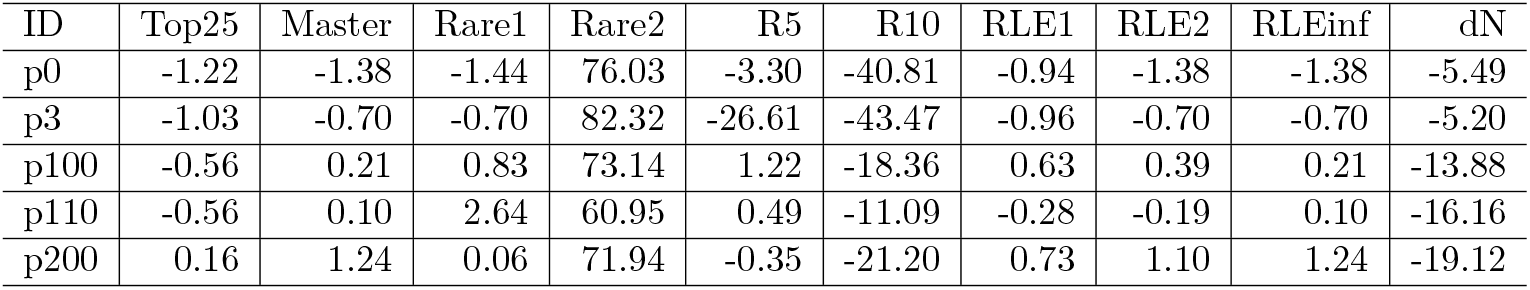
Relative differences in estimated standard deviations. Bootstrap versus analytic values (bootstrap-analytic)/analytic x 100.

| ID | Top25 | Master | Rare1 | Rare2 | R5 | R10 | RLE1 | RLE2 | RLEinf | dN |
| --- | --- | --- | --- | --- | --- | --- | --- | --- | --- | --- |
| p0 | -1.22 | -1.38 | -1.44 | 76.03 | -3.30 | -40.81 | -0.94 | -1.38 | -1.38 | -5.49 |
| p3 | -1.03 | -0.70 | -0.70 | 82.32 | -26.61 | -43.47 | -0.96 | -0.70 | -0.70 | -5.20 |
| p100 | -0.56 | 0.21 | 0.83 | 73.14 | 1.22 | -18.36 | 0.63 | 0.39 | 0.21 | -13.88 |
| p110 | -0.56 | 0.10 | 2.64 | 60.95 | 0.49 | -11.09 | -0.28 | -0.19 | 0.10 | -16.16 |
| p200 | 0.16 | 1.24 | 0.06 | 71.94 | -0.35 | -21.20 | 0.73 | 1.10 | 1.24 | -19.12 |

**Table 10:** Relative differences in estimated standard deviations. Jackknife versus analytic values as (jackknife-analytic)/analytic x 100.

| ID | Top25 | Master | Rare1 | Rare2 | R5 | R10 | RLE1 | RLE2 | RLEinf | dN |
| --- | --- | --- | --- | --- | --- | --- | --- | --- | --- | --- |
| p0 | -0.18 | -0.96 | -0.36 | 194.83 | 0.15 | -30.78 | 2.73 | -0.95 | -0.96 | 1.60 |
| p3 | -0.63 | 1.09 | 1.09 | 212.82 | -18.53 | -15.18 | 3.24 | 1.09 | 1.09 | 1.52 |
| p100 | 2.17 | -2.01 | 0.55 | 160.20 | 0.89 | -15.50 | 4.07 | -1.25 | -2.01 | -10.67 |
| p110 | 0.41 | -0.47 | 0.33 | 159.78 | 1.18 | -0.30 | 4.51 | -0.18 | -0.46 | -12.36 |
| p200 | -0.64 | 0.64 | 0.64 | 177.87 | -0.69 | -0.08 | 4.32 | 0.78 | 0.64 | -15.56 |

**Figure 1:**
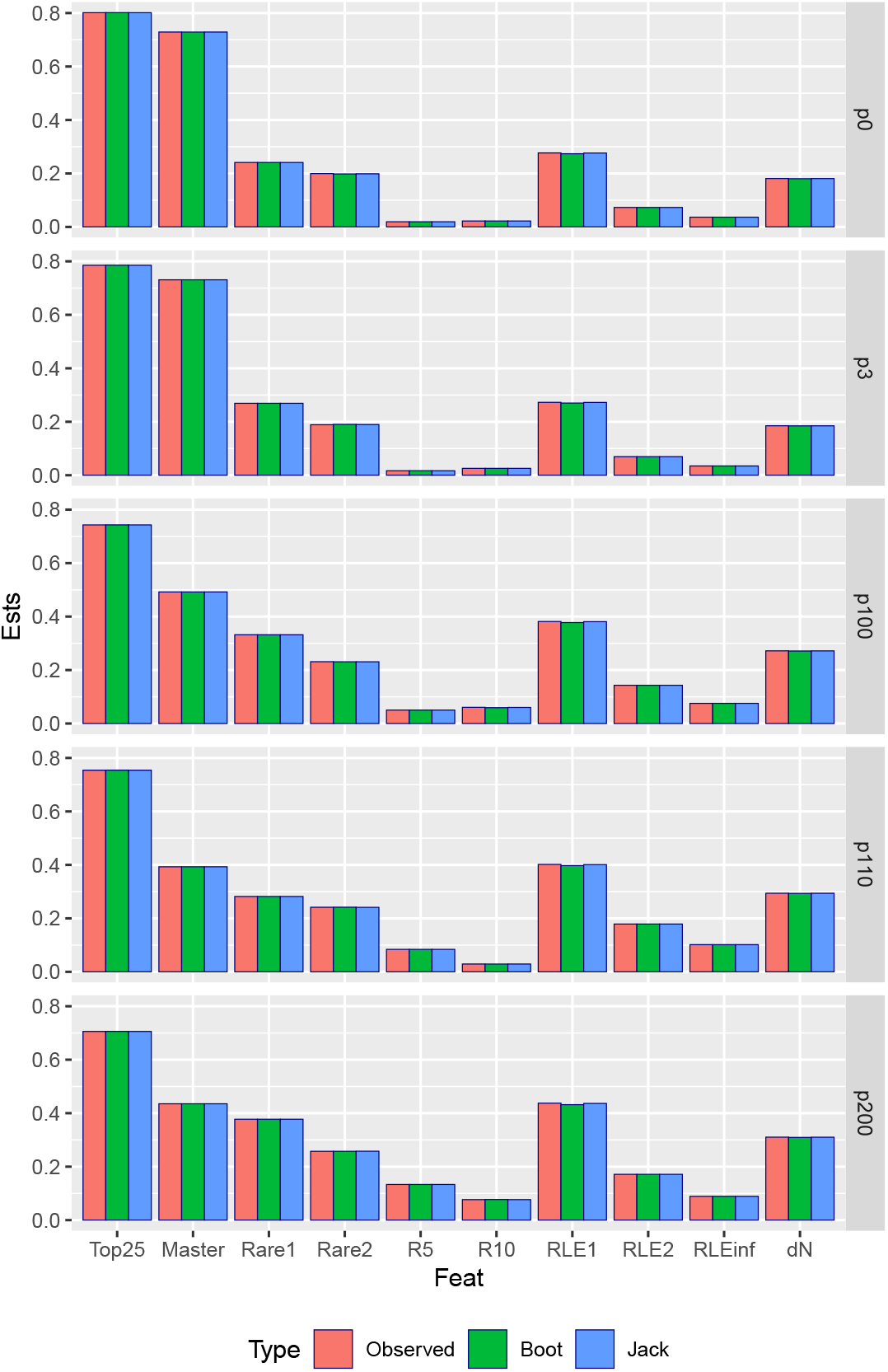
Observed and estimated indicator values by bootstrap and random delete-d jackknife

**Figure 2:**
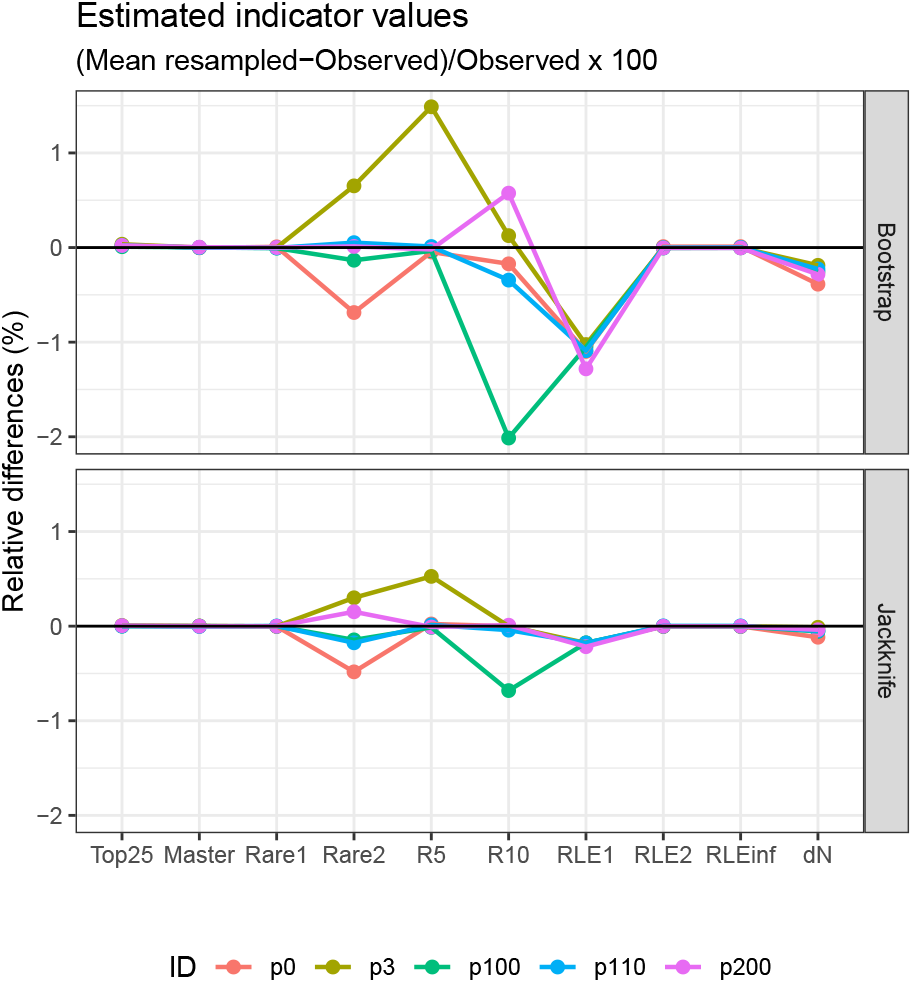
Relative differences in estimated indicator values as (Resampled-Observed)/Observed expressed as percentages.

Both resampling methods yield mean values closely matching those observed on the full sample for Top25, Master, and Rare1. Differences for *Rare2* remain below 1% with either method. The overall bias profiles of the bootstrap and jackknife are similar; however, bootstrap estimates display larger deviations from the observed values (Figure 2). Across all indicators and quasispecies, jackknife mean values deviate by less than 1%, whereas bootstrap estimates show larger biases by more than double.

*RLE*_1_ show consistent downward biases, whereas *Rare2* and *R*_10_ display sample-dependent biases with reference to the full sample observed values.

#### 3.5.2 Estimated standard deviations, resampled vs analytic

The standard deviations (SD) obtained analytically (Table 4) were compared with those estimated by bootstrap (Table 5) and random delete-*d* jackknife resampling (Table 6), each performed over 5000 iterations, as shown in Figure 3. Table 9 reports the relative differences between analytical and bootstrap SD values (in percentages), while Table 10 presents the corresponding relative differences between analytical and jackknife estimates.

**Figure 3:**
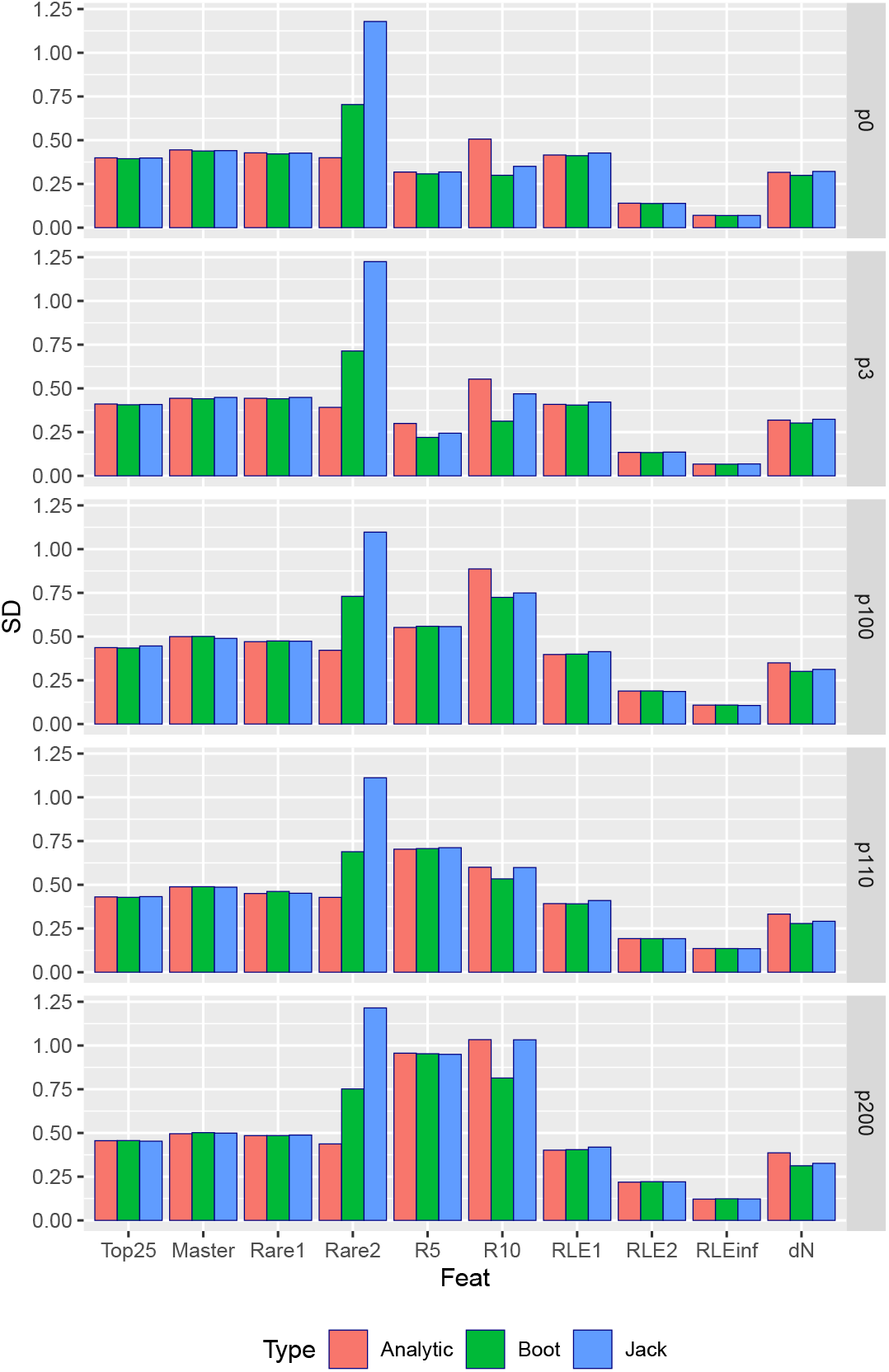
Analytic standard deviations and estimated by bootstrap and jackknife resampling methods.

*QFF* fractions show consistent values across the three variance estimation methods, except for *Rare2*, where notable differences emerge: bootstrap estimates are more than 50% higher than exact Bernouilli variances, and jackknife estimates are more than double of analytic values. For *RLE*_1_, bootstrap aligns more closely with analytic values (within 1%) than jackknife (within 2.7–4.5%). Both *RLE*_2_ and *RLE∞* exhibit similar trends, with bootstrap estimates within 1.5% and jackknife within 2% of the corresponding analytic values. The differences observed for the *R*_*k*_ indicators display diferences that are sample-dependent, and are challenging to interpret. *R*_5_ values generaly agree closely among all methods except for quasispecies HCV p3, whereas *R*_10_ shows significant differences but jackknife estimates better approximate analytical results.

### 3.6 Overdispersion and sensibility to rare variants

*QFF* fractions resampled by bootstrap and jackknife, as estimated proportions, are analyzed in the next two sections to investigate potential deviations from the expected resampling behavior, primarily driven by the high prevalence of haplotypes at very low frequencies.

A significant portion of reads in each quasispecies correspond to singletons—3.7% in HCV p0, 5.1% in HCV p100, and 7.0% in HCV p200. The cumulative fraction of reads for haplotypes with read multiplicity up to 10 increases to 6.0% in HCV p0, 9.4% in HCV p100, and 13.7% in HCV p200. For haplotypes with read multiplicity up to 100, these fractions rise further to 18.6%, 20.4%, and 24.5% respectively. Notably, the fractions approximately double with each increase in this multiplicity threshold (from 1 to 10, and from 10 to 100). Such a structure induces a heavy-tailed abundance distribution, where most mass is concentrated in a few abundant haplotypes but many extremely rare ones also contribute significantly. These features strongly affect resampling-based variance estimates because they may contribute to violate smoothness assumptions on which jackknife consistency depends, despite the delete-d resampling scheme, and introduce extra-binomial variation (overdispersion) in multinomial-like bootstrap sampling.

#### 3.6.1 Overdispersion in bootstrap resampling

Table 11 presents the results of the overdispersion analysis with p100 quasispecies, for each QFF indicator, including the following data: *p*, the observed fraction; *p*.*var*, the exact Bernoulli variance; *boot*.*var*, boostrap estimated variance corresponding, *ratio* the ratio of bootstrap estimated variance to the exact Bernouilli value; and *phi*, the overdispersion factor from the fitted Beta distribution to the bootstrap resampled values.

**Table 11:**
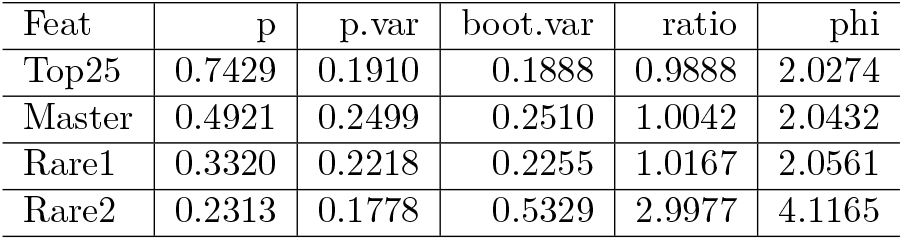
Bootstrap overdispersion factors. p100 quasispecies.

| Feat | p | p.var | boot.var | ratio | phi |
| --- | --- | --- | --- | --- | --- |
| Top25 | 0.7429 | 0.1910 | 0.1888 | 0.9888 | 2.0274 |
| Master | 0.4921 | 0.2499 | 0.2510 | 1.0042 | 2.0432 |
| Rare1 | 0.3320 | 0.2218 | 0.2255 | 1.0167 | 2.0561 |
| Rare2 | 0.2313 | 0.1778 | 0.5329 | 2.9977 | 4.1165 |

The overdispersion factors, *ϕ*, are close to 2 for *Top25, Master*, and *Rare1*, the fractions collecting the haplotypes with higher frequencies. *Rare2* with the fraction of reads for haplotypes with abundances ≤ 0.01% result in an overdispersion factor of 4. The bootstrap variances are very close to the exact Bernouilli variances for the three top indicators, but are the triple for *Rare2* (Table 11).

Note that *ϕ* is defined as the ratio of the observed variance to that expected under the sampling model, which in this context is multinomial for bootstrap resampling. Whereas the *ratio* in Table 11 shows in what extend the observed variance exceeds the Bernouilli value for the observed proportion in the full sample.

Bootstrap resampling with replacement effectively samples from a multinomial model, which reproduces relative frequencies according to observed probabilities. When the number of unique haplotypes is large and many of them appear only once, replacement resampling duplicates some rare observations, mimicking the stochasticity expected from independent multinomial draws. However, the notably high ratio of variances for *Rare2* between the bootstrap estimate and the exact Bernouilli value, roughly a factor of three, suggests that for heavy-tailed and sparse sets, multinomial sampling amplifies tail fluctuations.

As a result bootstrap resampling amplifies variability disproportionately when applied to rare variants that make up a large portion of the quasispecies at very low frequencies. The likely cause is the inherent limitation of bootstrap methods, where resampling with replacement results in some reads being duplicated while others remain unobserved (the 0.632 probability issue). Besides, it is important to note that although *Rare2* may be modeled as a binomial parameter, it actually represents an aggregate of a large number of non-independent multinomial parameters with non-zero covariances. The high dimensionality of this component group means that these covariances could substantially influence the overall variance estimated by resampling methods.

In summary, the bootstrap’s higher variance estimates for *Rare2* relative to the exact Bernouilli value likely arise from the bootstrap’s difficulty to accommodate the complex joint dependence of rare haplotypes despite the high number of iterations done, together with the 0.632 probability issue.

#### 3.6.2 Overdispersion in jackknife resampling

Table 12 presents the results of the overdispersion analysis on jackknife resampled values with HCV p100 quasispecies, for each *QFF* indicator, including the following data: *p*, the observed fraction; *p*.*var*, the exact Bernoulli variance; *jack*.*var*, variance estimated by the jackknife, *ratio* the ratio of jackknife estimated variance to the exact Bernouilli value; and *phi*, the overdispersion factor from the fitted Beta distribution to the jackknife resampled values.

**Table 12:**
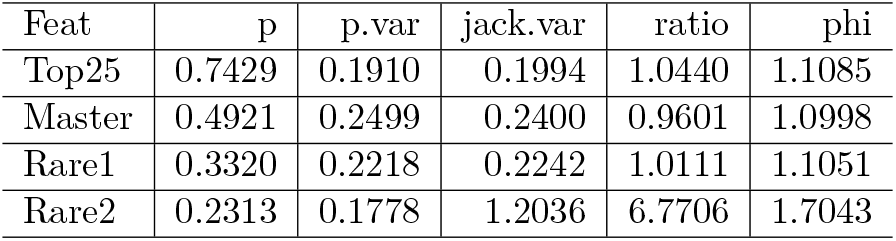
Jackknife overdispersion factors. p100 quasispecies.

The overdispersion factor, *ϕ*, is close to 1 for *Top25, Master*, and *Rare1*, indicating that sampling from a hypergeometric distribution behaves well for these indicators. In contrast, *Rare2* displays poorer behavior, with *ϕ* = 1.7. These results indicate that resampling aligns better with the hypergeometric model than with the multinomial model showing higher *ϕ* values. However, the ratio of observed variance to the exact value remains near 1 for the top three indicators, but rises dramatically to 6.8 for *Rare2*.

Similar values were obtained for the other quasispecies (data not shown).

The results are counterintuitive in that the *a priori* superior resampling model based on the hypergeometric distribution yields poorer performance than the bootstrap’s multinomial resampling.

The classical jackknife’s stringent functional smoothness requirements are relaxed under the delete-d formulation. By removing multiple observations per replicate, the delete-d jackknife averages the deletion effects across several data points rather than a single one, thereby enhancing robustness for non-smooth or discontinuous statistics. According to asymptotic results by Shao and Wu [19–22], selecting *d* such that 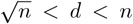 ensures stable variance and mitigates bias for estimators that deviate from smoothness. In the present analysis, with *n* ≈ 100 000, the lower bound 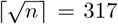 indicates that deleting 10% of the sample—far exceeding this threshold—comfortably satisfies the conditions for consistent and approximately unbiased variance estimation. However, for heavy-tailed non-smooth statistics dominated by rare categories in complex quasispecies, such as *Rare2*, this fraction may be insufficient to completely dominate the lack of smoothness. This is explored in the next section with deleted fractions ranging from 5 to 70%.

### 3.7 Sensibility of jackknife to the deleted fraction *d*

The fraction of reads removed in the jackknife was selected *a priori* to be 10%, as this proportion was deemed sufficient to introduce meaningful perturbations in the haplotype distribution of the resamples, while still minimizing bias relative to the full quasispecies sample. To evaluate the effect of the delete fraction on the estimated variances, several values of *d* were examined: 5%, 10%, 20%, 30%, 50%, and 70% each with 5000 resampling iterations.

Figure 4 shows the relative differences in mean values at each deleted fraction compared to the values observed on the full sample for quasispecies HCV p0, illustrating the sensitivity of each indicator to the fraction deleted in the jackknife. Correspondingly, Figure 5 displays the estimated standard deviations for these indicators. Equivalent results for quasispecies HCV p100 are presented in Figures 6 and 7. The HCV p0 quasispecies represents a population of low to moderate diversity, whereas HCV p100 corresponds to a quasispecies matured over 100 passages in a non-coevolving environment, showing a 2.2-fold fitness increase relative to HCV p0 [45,46], and with higher levels of diversity and evenness.

**Figure 4:**
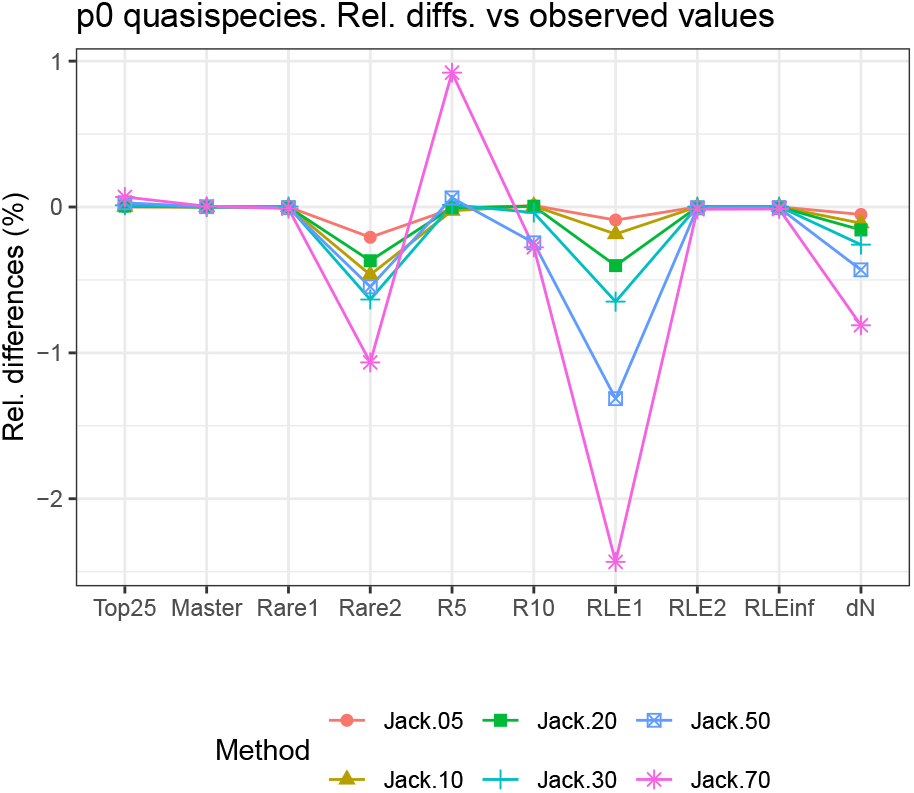
HCV p0 quasispecies. Relative differences expresed in percentage between values observed on the full sample and jackknife mean values obtained at different deleted fractions, as (obs - jack)/obs x 100.

**Figure 5:**
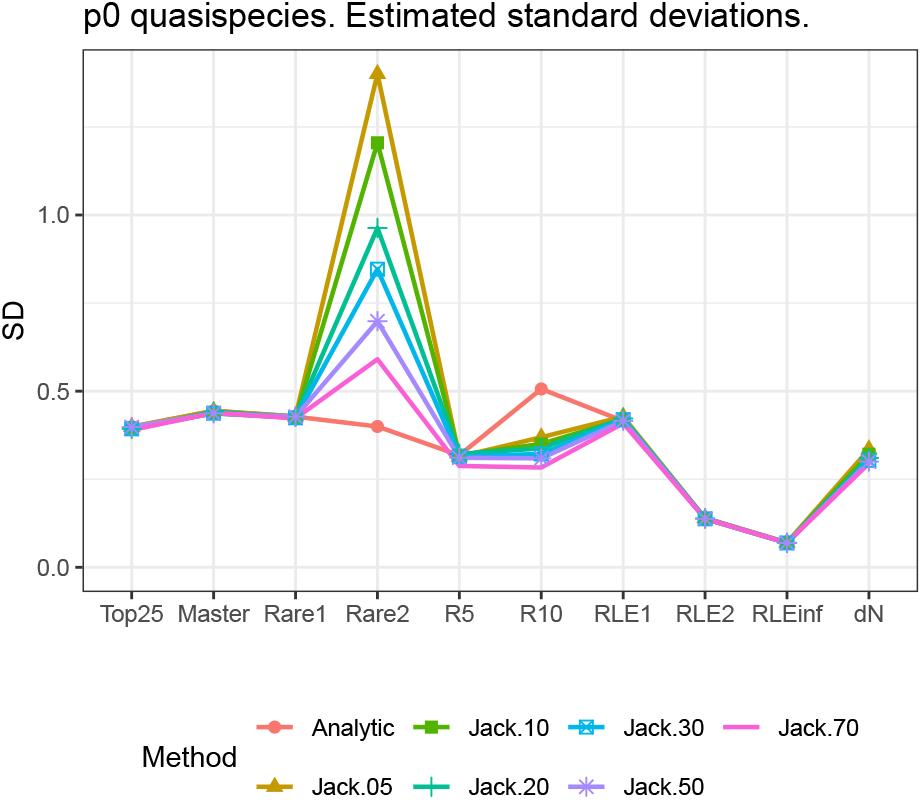
HCV p0 quasispecies. Analytic and the random delete-d jackknife estimated standard deviatios at different *d* values, from 5% to 70%.

**Figure 6:**
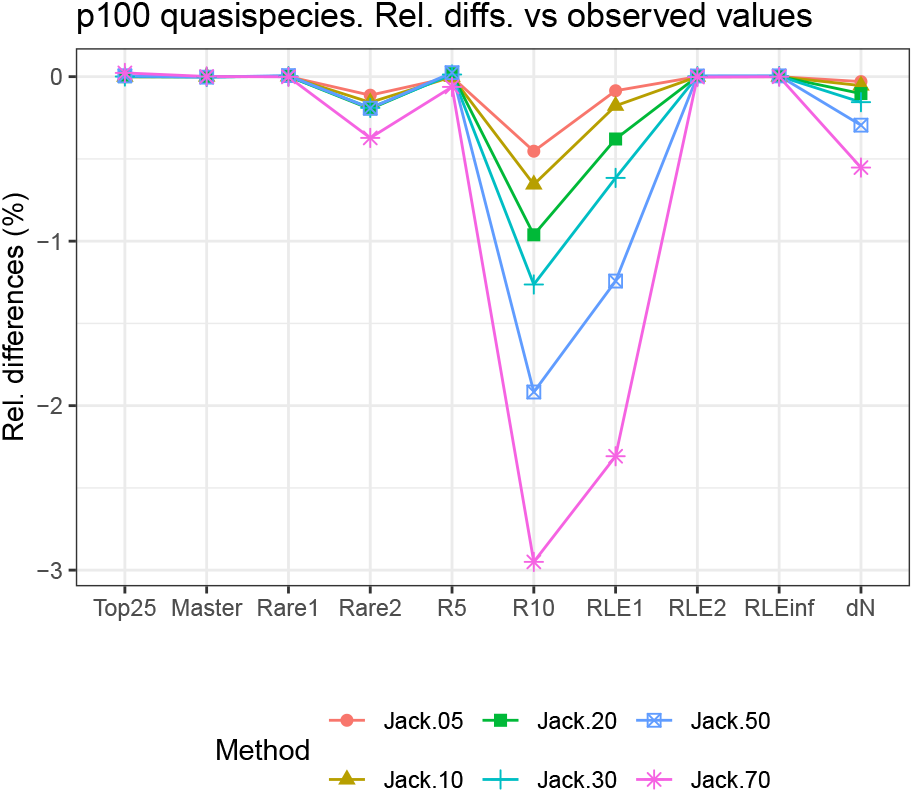
HCV p100 quasispecies. Relative differences expresed in percentage between values observed on the full sample and jackknife mean values obtained at different deleted fractions, as (obs - jack)/obs x 100.

**Figure 7:**
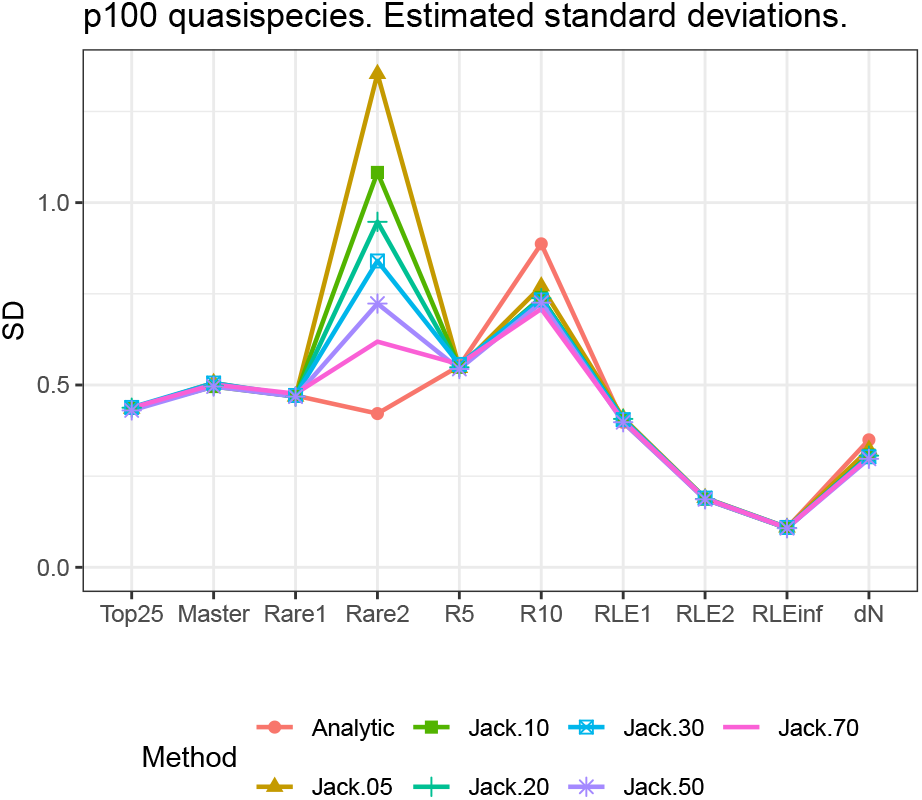
HCV p100 quasispecies. Analytic and the random delete-d jackknife estimated standard deviatios at different *d* values, from 5% to 70%.

In both quasispecies, *Master, Top25, Rare1, RLE*_2_ and *RLE∞* are robustly estimated at all deleted fractions. *Rare2* accuses moderated increasing biases (below 1% for p0, below 0.5% for p100) at larger deleted fractions. The indicators *R*_10_ and *RLE*_1_ are increasingly sensitive to larger deleted fractions, with moderated differences below 3%. However estimated standard deviations for *RLE*_1_ are robust to these changes. While the bias in jackknife estimates of standard deviations for *Rare2* progressively corrects as the deleted fraction increases, at 70% deletion it still exhibits noticeable deviation. Conversely, standard deviation estimates for *R*_10_ remain largely unaffected by increasing deleted fractions maintaining its bias with reference to the analytic values.

Seemingly, the lower bound on *d* given by the asymptotic theory of Shao and Wu [19], expressed as 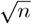, proves largely insufficient for complex, highly skewed populations and massive sample sizes, as in our case, where even deleting 70% of the sample fails to solve smoothness issues with *Rare2*.

### 3.8 Bootstrap and Jackknife with H random

Standard deviation estimates for *RLE*_*q*_ indicators were obtained using both bootstrap and random delete-10% jackknife under two scenarios: (i) with *H* treated as known and fixed, taken from the haplotype count in the full sample, and (ii) with *H* treated as random, corresponding to the haplotype cardinality in each resample. Figure 8 compares these results with the standard deviations derived analytically using the delta method, and Figure 9 shows the relative differences between the estimated and analytic standard deviations.

**Figure 8:**
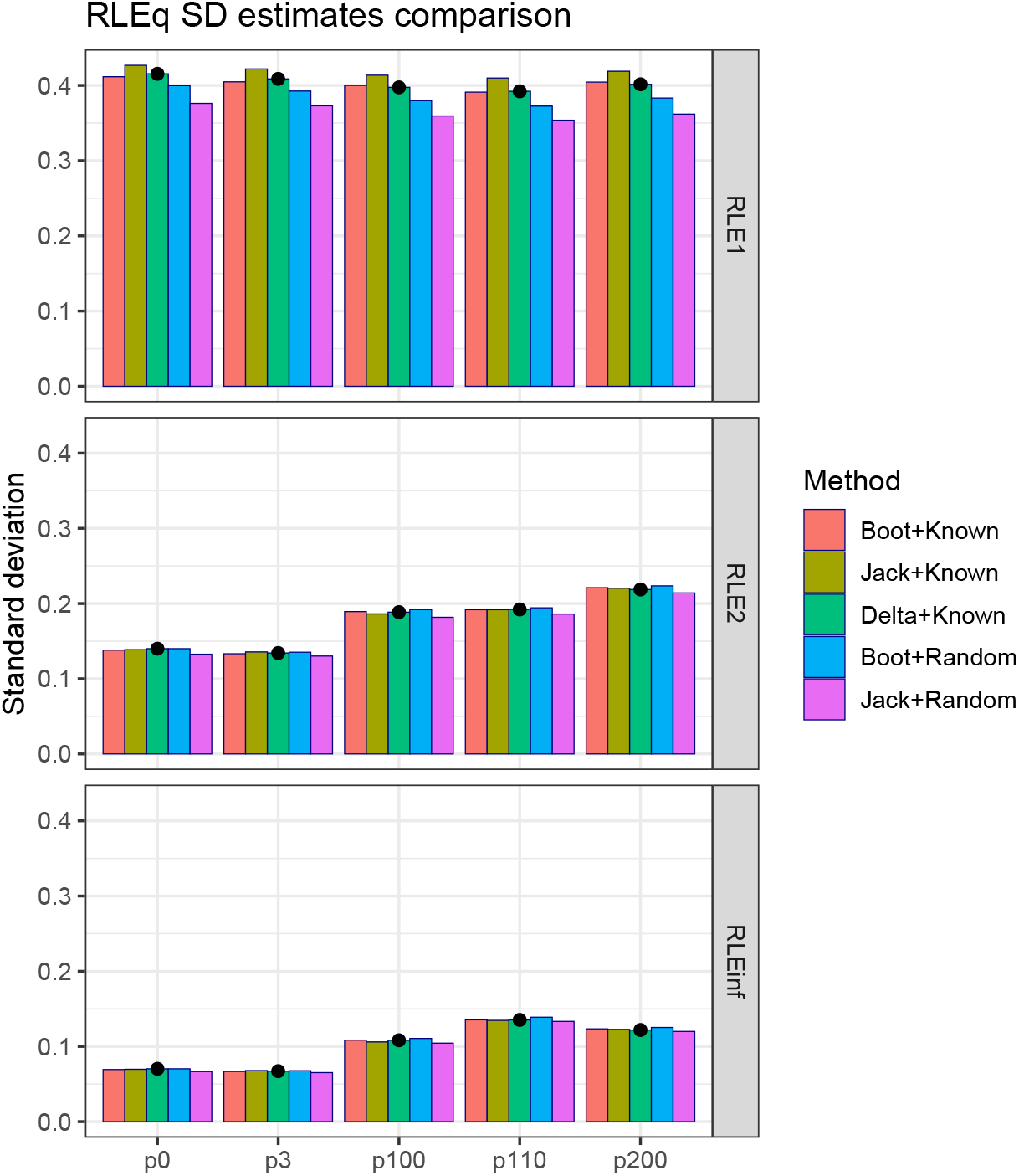
Comparison of RLE standard deviations by the methods: Delta, bootstrap + H known, jackknife + H known, bootstrap + H estimated, and jackknife + H estimated. Black dot on the analytic value.

**Figure 9:**
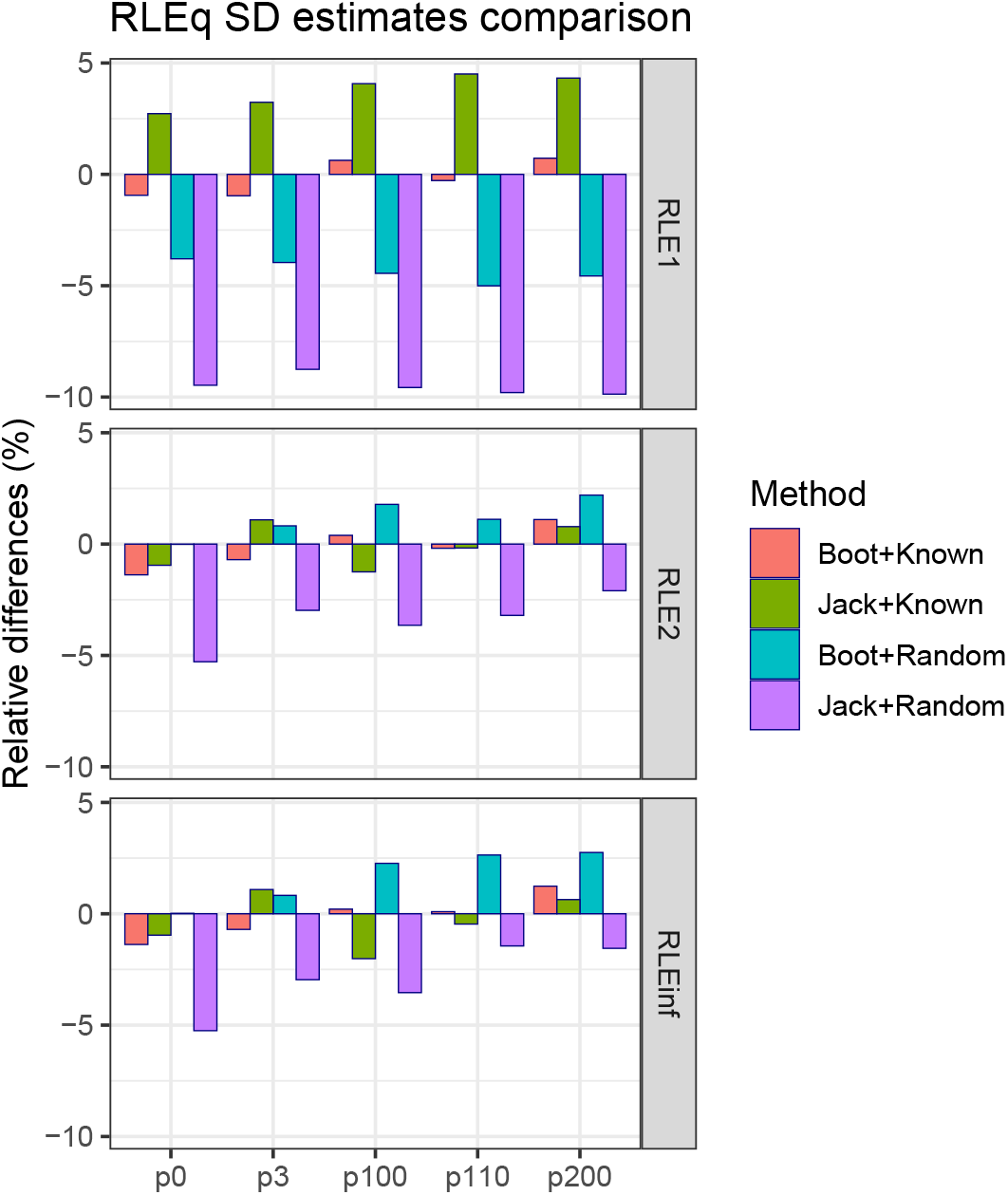
Relative differences in RLEq standard deviations with respect to the delta method values for the following methods: bootstrap with H fixed, jackknife with H fixed, bootstrap with H estimated, and jackknife with H estimated. Expressed as (resampled-analytic)/analytic x 100.

Among the four resamling methods, the bootstrap with fixed *H* yields most similar values to the analytic estimates, for the three *RLE*_*q*_ indicators; whereas the jackknife with random *H* shows the greatest divergence.

For *RLE*_1_, the bootstrap with fixed *H* yields standard deviation values close to the analytic estimates, whereas the jackknife with fixed *H* consistently produces higher values. Under the random *H* scenario, both bootstrap and jackknife produce lower values than the analytic ones, with the jackknife showing the largest underestimation. For *RLE*_2_ and *RLE*_*∞*_, the differences among the three methods are smaller than those observed for *RLE*_1_; however, the jackknife with random *H* still yields consistently lower standard deviations; by contrast the bootstrap with random *H* yields consistently higher standard deviations.

*RLE*_*q*_ indicators are expressed as the ratio of the logarithm of Hill numbers of order *q* to the logarithm of the number of haplotypes. *RLE*_1_ is a function of the sum of the *p*_*i*_ {ln(*p*_*i*_)} terms, where the ln(*p*_*i*_) factor increases the sensitivity to low-frequency haplotypes. In contrast, *RLE*_2_ is a function of the sum of 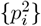 terms, which reduces the influence of rare haplotypes. *RLE* depends solely of the frequency of the master haplotype, the most robust in resampling. The differing dependency of *RLE*_*q*_ indicators on rare haplotypes helps explain why the discrepancies among standard deviation estimates follow the order *RLE*_1_ *> RLE*_2_ *> RLE*_*∞*_, with *RLE*_1_ showing the largest differences.

## 4 Discussion

Variance estimation for quasispecies indicators based on single NGS observations presents intrinsic challenges due to the absence of replicates and the highly skewed distributions typical of RNA viral mutant spectra. In the context of the selected quasispecies indicators, except for exact variances for QFF fractions, derived from the Bernoulli formula, no variance estimation method is free of limitations. Nevertheless, functionals such as *RLE*_*q*_ and *R*_*k*_ require approximate methods.

The delta method assumes local linearity through a first-order Taylor expansion, which fails for highly nonlinear or discontinuous statistics. However, expanding to the second term does not resolve potential inconsistencies—if the function is not sufficiently smooth or if higher derivatives are large, higher-order terms may lead to overestimation o inaccuracies, especially when the function’s curvature varies greatly. Bootstrap resampling is affected by the 0.632 bias and inflated variance under heavy-tailed abundance distributions. The jackknife demands smoothness of the statistic, an assumption often violated in viral quasispecies dominated by rare haplotypes. The inherent skewness of quasispecies distributions limits the precision of both resampling schemes.

Across all studied quasispecies, variance estimates for the fractions *Master, Top25*, and *Rare1* obtained by resampling methods consistently match the exact Bernouilli values. Whereas resampled variances for *RLE*_*q*_ indicators match the values calculated with analytic expressions, under the *H* known (not-random) hypothesis. The variance of *R*_5_ is also reliably estimated by all three approaches. In contrast, *Rare2*, and *R*_10_ indicators exhibit notable discrepancies between analytic and resampling-based estimates.

As shown in Tables 9 and 10, and Figure 3 bootstrap-derived standard deviations for *RLE*_1_ indicators most closely align with analytical results-within 1%-while jackknife estimates can deviate by up to 4.5%. *RLE*_1_ jackknife-based standard deviations are all larger than the analytic counterpart. *RLE*_2_ and *RLE*_*∞*_ are equally estimated with both resampling methods, within 2% of the analytic values. Conversely, substantial differences are observed for *Rare2* and *R*_10_ between analytic and resampled variance estimates (Figure 3).

Very large sample sizes and highly skewed distributions complicate accurate variance assessment under these methods. In this context, the delta method offers an appealing analytic route because it exploits asymptotic normality of multinomial estimators to deliver closed-form approximations of variance for smooth statistics. Nevertheless, its accuracy ultimately depends on the local differentiability and linearity of the functional with respect to haplotype frequencies. These conditions may be compromised for measures involving logarithmic or rank-based transformations of rare haplotypes

Bootstrap resampling amplifies the impact of rare haplotypes through the 0.632 sampling imbalance, introducing moderate positive bias in mean indicator estimates (Table 7) and inflated variances for heavily tailed distributions such as *Rare2*. The deviation of the estimated *Rare2* SD under bootstrap with reference to exact Bernouilli value, with *Rare2* an appreciable fraction across all studied quasispecies (Table 3) - 0.189 to 0.294 - demostrates the limitations of the bootstrap for this kind of estimators with complex samples.

The random delete-*d* jackknife, which draws subsamples without replacement, displays complementary behavior: it recovers mean indicator values more accurately than the bootstrap (Tables 7 and 8, and Figure 2), but tends to overestimate the variance of *Rare2* and *R*_10_. Increasing the deleted fraction *d* reduces variance bias for *Rare2*, but at the expense of systematic shifts in mean values and instability for indicators dominated by rare haplotypes. This suggests that the relaxed smoothness requirements of delete-*d* jackknife compared to delete-1 do not fully accommodate the sparsity and high dimensionality of viral quasispecies data.

For *R*_5_, the agreement between delta, bootstrap, and jackknife estimates across all quasispecies supports the validity of the analytical approach for top-haplotype evenness metrics. This result mirrors what is observed for indicators such as *Top25* and *Master*, where dominant variants drive the signal and functional dependence on frequencies is approximately linear. Exceptions arise, as for HCV p3, where degeneracy near rank 5 introduces near-ties that disrupt indicator smoothness—likely explaining similar issues with *R*_10_.

Under the alternative random-*H* resampling scheme, *RLE*_*q*_ indicators exhibit systematic bias in their resampled mean values relative to observed full-sample values, stemming from a consistent underestimation of haplotype number in resamples—principally with the bootstrap due to the 0.632 problem, but also with the jackknife caused by the deleted fraction. Because *RLE*_*q*_ is normalized by by the log of the haplotype count, this bias leads to upwardly biased resampled estimates and propagates to standard deviation estimation.

Overall, the bootstrap provides an empirical benchmark that closely approximates analytic results in the presence of moderate nonlinearity, while delete-*d* jackknife improves mean recovery at the expense of variance stability when smoothness is weak. Among all methods compared, the first-order delta approximation stands out as the most reliable and efficient option for large-sample quasispecies analysis.

## 5 Conclusions

Collectively, these results support the first-order delta approximation as the most reliable and computationally efficient method for variance and standard error estimation in large-sample quasispecies analyses, particularly for functionals such as *RLE*_*q*_ and *R*_*k*_. For inference-based studies, binomial standard errors derived from observed proportions in the full sample remain appropriate for QFF indicators, while the delta method provides accurate and analytically justified approximations for complex functionals. Hypothesis testing using z-statistics and corresponding p-values is mathematically and empirically validated under these conditions. To complement significance testing, standardized effect sizes (e.g., Cohen’s *d*) should also be reported to quantify the magnitude of differences between quasispecies populations, providing biological context beyond statistical significance—especially in high-depth NGS datasets where sample sizes are inherently large. The pooled standard deviation required for effect size computation can be derived directly from estimated standard errors and sample sizes, ensuring methodological consistency. Together, these analytical and inferential tools establish a robust and interpretable statistical framework for assessing quasispecies diversity, effectively balancing computational efficiency, statistical rigor, and biological insight.

